# Human mitochondrial DNA variants influence telomere length: evidence from a transmitochondrial cybrid model

**DOI:** 10.1101/2025.06.27.661964

**Authors:** Manon Mahieu, Jean-Philippe Defour, Barbara Mathieu, Elena Richiardone, Isaac Heremans, Elisa Fabiole, Gabriel Levy, Gabriel Le Berre, Isabelle Scheers, Bénédicte Brichard, Thierry Arnould, Patrick Revy, Guido Bommer, Bernard Gallez, Cyril Corbet, Anabelle Decottignies

**Affiliations:** Genetic & Epigenetic Alterations of Genomes, de Duve Institute, UCLouvain, Brussels, Belgium; Cliniques Universitaires Saint-Luc, UCLouvain, Brussels, Belgium; CHC Liège, Liège, Belgium; Biomedical Magnetic Resonance Unit, Louvain Drug Research Institute, UCLouvain, 1200 Brussels, Belgium; Pole of Pharmacology and Therapeutics, Institut de Recherche Expérimentale et Clinique, UCLouvain, Brussels, Belgium; Biochemistry, de Duve Institute, UCLouvain, Brussels, Belgium; Laboratory of Genome Dynamics in Human Diseases, Equipe Labellisée Ligue 2026, INSERM UMR 1163, Imagine Institute, Paris, France; Université Paris Cité, Imagine Institute, Paris, France; Pediatric gastroenterology and hepatology unit, Cliniques universitaires Saint-Luc, Université catholique de Louvain, Brussels, Belgium; Laboratory of Biochemistry and Cell Biology, NAmur Research Institute for LIfe Sciences (NARILIS), UNamur, Namur, Belgium; WEL Research Institute, Wavre, Belgium

**Keywords:** Mitochondrial genome, Telomeres, Telomere length inheritance, Oxidative stress, Complex I

## Abstract

Telomere shortening is a hallmark of aging, yet telomere length (TL) varies considerably among individuals and is strongly influenced by inheritance. In mice, efficient mitochondrial function-characterized by low reactive oxygen species (ROS) production-is critical for telomere elongation during early embryogenesis. Since mitochondrial DNA (mtDNA) encodes several subunits of the electron transport chain, it may influence TL at birth by regulating mitochondrial function *in utero*. To explore the relationship between mtDNA and TL in human, we used a transmitochondrial cybrid approach, introducing mitochondria from donor platelets with varying telomere lengths into mtDNA-depleted cells. This revealed an inverse correlation between donor blood TL and mitochondrial ROS levels measured in the resulting cybrids, suggesting that specific mtDNA variants may contribute to the maintenance of long telomeres in humans by enhancing mitochondrial fitness. During *in vitro* cybrid formation, a transient phase of oxidative stress precedes cellular adaptation. In this specific window, mtDNA variants associated with reduced complex I (CI) activity induced rapid telomere shortening—an effect rescued by antioxidant and NAD⁺ precursor supplementation. While these variants occur naturally in certain individuals with long telomeres, our data suggest that, at least under *in vitro* conditions of acute oxidative stress, CI is critical to support PARP1 activity by maintaining the NAD⁺/NADH balance, thereby preserving telomere integrity. Collectively, these findings solidify the link between mtDNA variants and human TL regulation, highlighting potential therapeutic opportunities for mitochondrial replacement strategies.

**Significance Statement:** Telomere length at birth influences aging trajectories and disease risk later in life, yet the mechanisms governing this trait remain incompletely understood. Using a transmitochondrial cybrid approach, we show that single-nucleotide variants in the mitochondrial genome of healthy donors directly affect mitochondrial metabolism and reactive oxygen species production. In addition, mitochondrial ROS levels measured in cybrids inversely correlate with blood cell telomere length in donors. During cybrid formation, mitochondrial DNA variants associated with reduced CI activity promote telomere shortening. Attrition was reversed by antioxidant and NAD⁺ precursor supplementation, pointing to an essential role for robust CI function in sustaining telomere length during acute oxidative stress, at least under *in vitro* conditions. Together, these findings establish a direct link between mitochondrial genetics, redox homeostasis, and telomere maintenance in human cells.

## Introduction

Telomeres form protective caps at chromosome ends thanks to the binding of a six-member protein complex termed shelterin (1). With cell divisions, telomeres of normal somatic cells progressively shorten, a process that likely contributes to cellular aging and the development of various age-related pathologies (2,3). Accordingly, cohort studies established that shorter leucocyte telomere length (TL) was associated with a higher risk of diseases affecting specific organs, such as the lung or the liver, and an increased incidence of esophageal cancer, as well as lymphoid and myeloid leukemia (4). However, some pathogenic mutations in *POT1* gene, linked to very long telomeres, were also previously associated with increased risk for lymphoid and myeloproliferative neoplasms (5,6), underscoring the complex relationship between TL and cancer predisposition.

While aging is a key determinant of TL, genetic variation also plays a significant role in the variability of leukocyte TL among healthy individuals, as evidenced by the identification of nuclear genomic SNPs associated with TL (7). Heritability estimates for human TL reach 82% (8). Consistently, TL in the zygote appears to play a key role in determining TL later in life (7), indicating that it is largely established during early embryonic development. Telomere length resetting during embryogenesis is a telomerase-dependent process that occurs at the morula-to-blastocyst transition (9). Recent findings in mice show that experimentally disrupting mitochondrial function in zygotes significantly impairs telomere elongation between the 8-cell and blastocyst stages, ultimately leading to shorter telomeres in the offspring (10). Similar outcomes have been observed in offspring from reproductively aged or obese female mice-two conditions known to compromise oocyte mitochondrial integrity (10). Collectively, these findings underscore the critical role of proper mitochondrial function in telomere regulation during embryogenesis. Further evidence comes from individuals with mitochondrial disorders such as MELAS syndrome and Leber’s hereditary optic neuropathy (LHON). These individuals exhibit significantly shorter leukocyte telomeres at birth, likely reflecting impaired TL establishment during early development (11).

While damaged mitochondria are associated with shorter TL in mouse embryos and may impair TL regulation in patients with mitochondrial diseases (11), the impact of naturally occurring mitochondrial genome SNPs on TL remains largely unexplored. With a total of 29 haplogroups and a myriad of subhaplogroups characterized by distinct variants compiled in the Mitomap database (12), human mtDNA genomes display large heterogeneity. Some variants within the coding regions of mtDNA, such as the NADH dehydrogenase subunits (ND1, 2, 3, 4, 4L and 5) of complex I (CI), Cytochrome b, the COXI, II and III subunits of CIV or the ATP6 and 8 subunits of the ATP synthase (CV) lead to distinct electron transport chain (ETC) properties that impact mitochondrial metabolism and ROS production. In line with the hypothesis that the mitochondrial genome may contribute to the observed stronger maternal influence on TL at birth (8,13)-given that mtDNA is maternally inherited-, we hypothesized that non-pathogenic variation in the mtDNA sequence could affect TL.

Building on reference curves for TL in the Belgian population, obtained through fluorescence *in situ* hybridization (FISH) combined with flow cytometry (Flow-FISH), we identified mtDNA variants associated with either very long or short leukocyte TL. We leveraged a transmitochondrial cybrid approach to directly test the effects of these mtDNA variants on mitochondrial function in mtDNA-depleted recipient cells and uncovered an inverse relationship between leukocyte TL in the donors and mitochondrial ROS levels measured in the cybrids. Our results further highlight the critical role of mtDNA-encoded CI activity in sustaining NAD⁺ metabolism and telomere integrity, particularly *in vitro*, under oxidative stress conditions arising during the metabolic transition from glycolysis to oxidative phosphorylation in early cybrid development. Collectively, these findings reinforce the link between mtDNA-encoded mitochondrial function and telomere regulation in human cells.

## Results

### Establishment of reference curves for TL using Flow-FISH

To establish reference curves for TL in Belgium, we applied the gold standard Flow-FISH approach (14) on leukocytes isolated from a cohort of 491 healthy donors aged 7 days to 99 years old. Percentile curves for TL in lymphocytes (Table S1 and Fig 1A) and granulocytes (Table S1 and Fig 1B) were comparable to those obtained previously (15,16) (Fig S1A) and showed large heterogeneity among individuals of similar age. Although still controversial, obesity may negatively impact TL (7,17). In our cohort, average BMI was not statistically different between donors with telomeres below the 5^th^ or above the 90^th^ percentile (Fig S1B). Given previous evidence supporting a stronger maternal inheritance of TL in humans (8,13), with similar trend observed in birds (18), we sought to investigate the potential contribution of the human mitochondrial genome to TL heterogeneity. To this end, we selected seven individuals with long, average or short telomeres from our Flow-FISH curves (Fig 1C). All were of European origin, thus excluding a possible bias linked to African ethnic groups (7), and the seven donors belonged to distinct mitochondrial subhaplogroups (Fig 1D). All but one (#5) were female donors. Strikingly, donor #6, with lymphocyte telomeres at the 95^th^ percentile, turned out to belong to the K1a subhaplogroup that encompasses the centenarian-enriched A177T variant (m.G9055>A) in the ATP6 subunit of ATP synthase (CV) (19,20) with an allele frequency of 5.4% in Mitomap database (Fig 1D).

**Figure 1.**
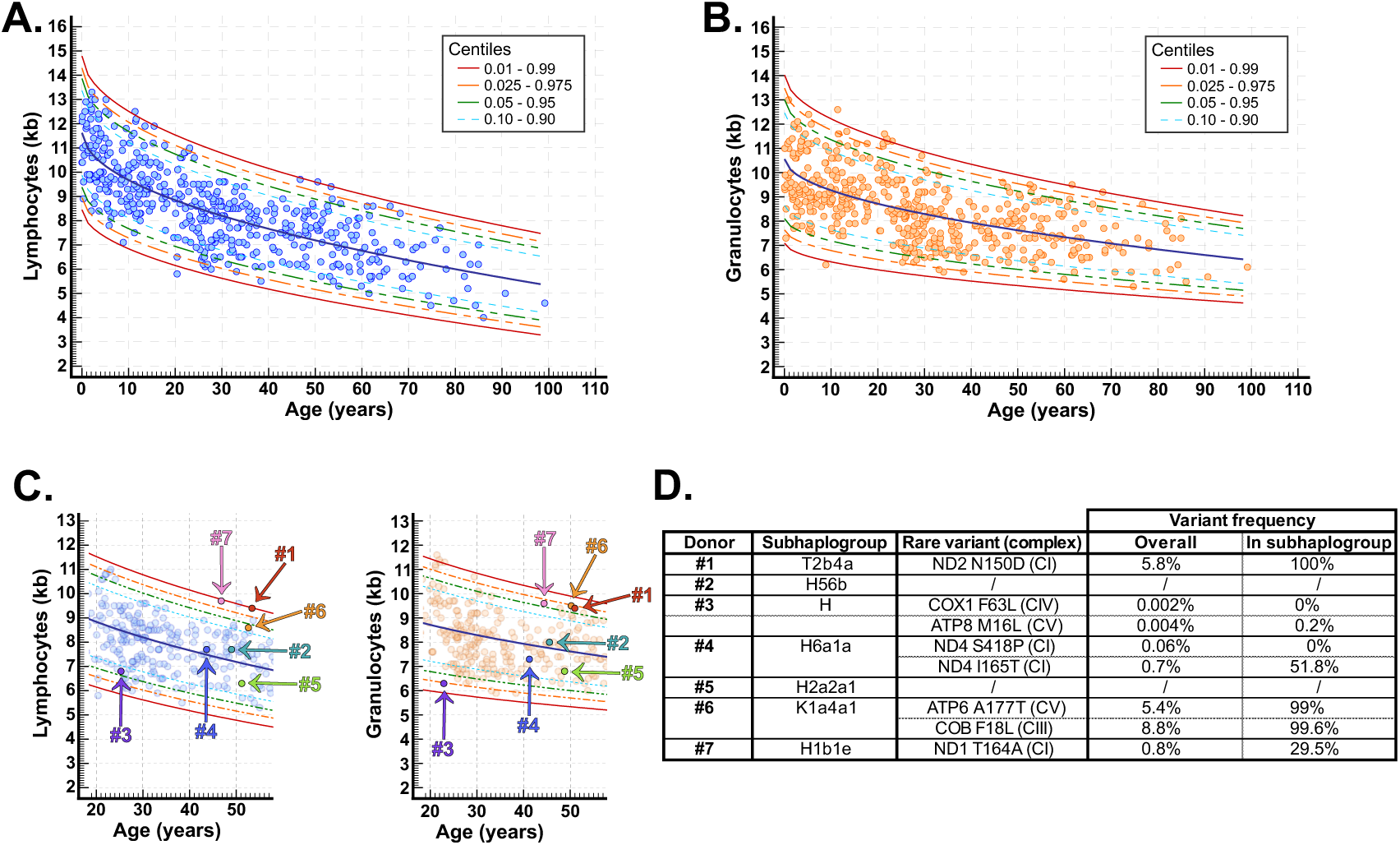
Establishment of Flow-FISH reference curves for telomere length A. The age-related reference interval and centiles were calculated after telomere length measurement by Flow-FISH in lymphocytes obtained from 491 healthy donors. The P50 curve is shown in dark blue. Other percentile curves are shown as indicated in the legend. **B.** Same as A in granulocytes. **C.** TL in lymphocytes (left) or granulocytes (right) of seven selected donors. **D.** Mitochondrial subhaplogroups of the seven selected donors with an overview of the variants showing both a change in the amino acid sequence and an overall frequency of less than 10% in Mitomap database in the mtDNA coding regions of the donors. The variant frequency in the subhaplogroup of each donor is indicated in the last column.

### Distinct mitochondrial genomes differently modulate TL in transmitochondrial cybrids

To evaluate the impact of distinct mitochondrial genomes on TL in human cells, we used a transmitochondrial cybrid strategy in which mtDNA-depleted Rho^0^ (ρ^0^) cells, derived from 143B human osteosarcoma cancer cells, are fused with donor platelets, thereby restoring functional mitochondria carrying donor-specific mtDNA variants (21). The presence of variants in cybrids was checked by sequencing and mtDNA content was evaluated by qPCR (Fig S2). Southern blot analysis revealed that telomeres in 143B ρ^0^ cells were shorter than in parental 143B cells (Fig 2A). To rule out clonal artifacts, mtDNA depletion was independently repeated, and multiple ρ^0^ clones were isolated, all of which exhibited a consistent reduction in TL (Fig S3A). This telomere shortening was not associated with decreased expression of the telomerase subunits h*TR* or h*TERT* (Fig S3B), but correlated with a reduction in telomerase activity measured *in vitro* using cell extracts (Fig S3C). In line with a role for mitochondrial function in supporting telomerase activity, fusion of 143B ρ^0^ cells with platelets from donors #6 and #7 (Cyb6 and 7) restored telomerase activity and rescued TL in ρ^0^ cells to levels comparable to those of parental 143B cells (Fig 2A-B and Fig S3D). Telomere elongation in these cybrids was accompanied by a reduced frequency of short telomeres, as assessed by TeSLA (Fig S3E). Unexpectedly, TL was not restored in Cyb1-5 cybrids. Instead, fusion with platelets from donors #1 and #2 induced a marked and reproducible telomere shortening in ρ^0^ recipient cells (Fig 2A and Fig S3D). This effect was not associated with reduced telomerase activity or decreased h*TR* or h*TERT* transcript levels, but rather with pronounced telomeric damage and telomere fusions following cybrid formation (Fig 2B-E).

**Figure 2.**
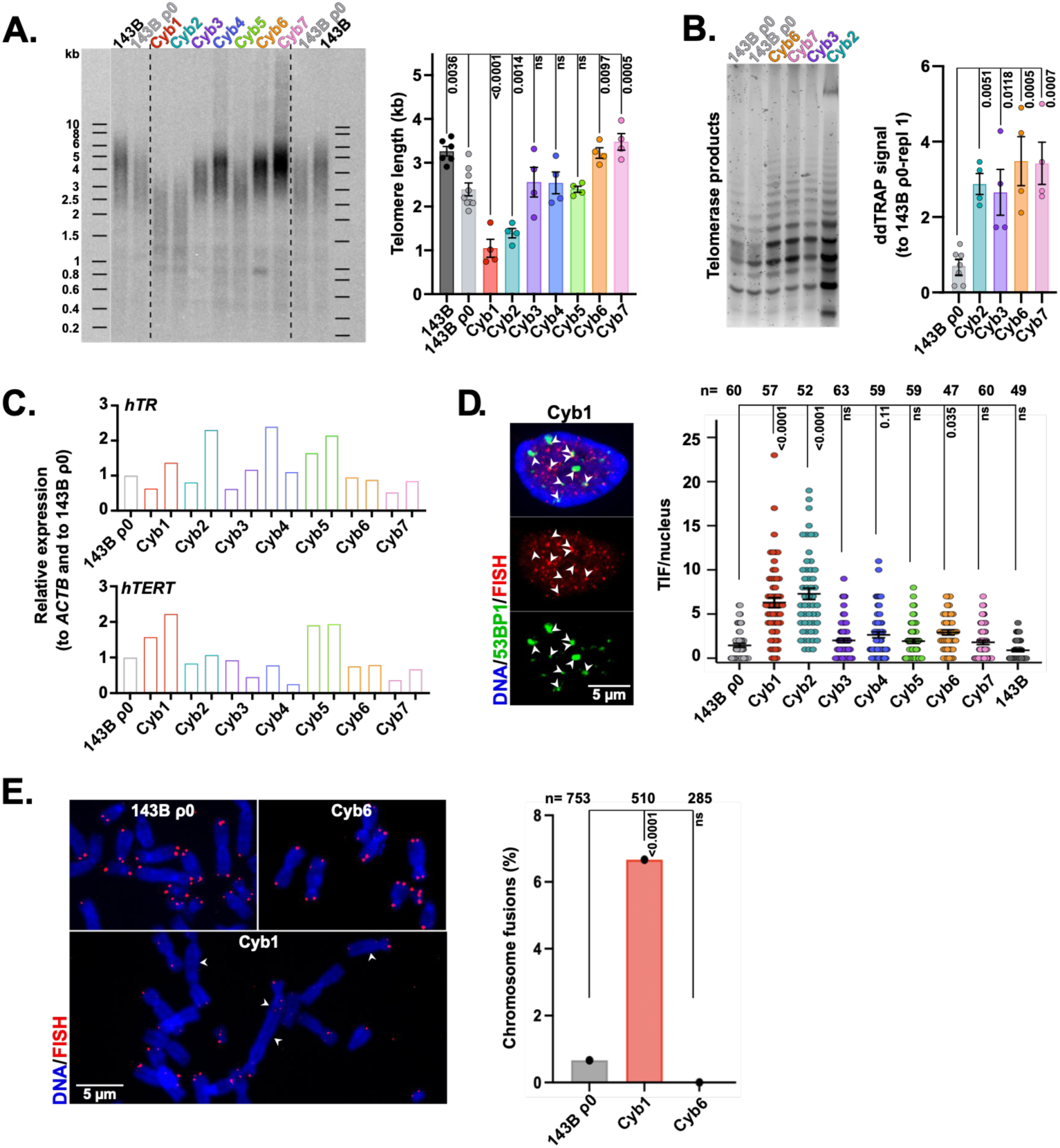
Mitochondrial genomes modulate telomere length and integrity of ρ^0^ recipient cells. **A.** Left: Representative Southern blot analysis (TRF) of telomeres in 143B, 143B ρ^0^ recipient cells and in one cybrid clone from each donor at the first PDs post-selection. Right: Quantification of TRF on four independent cybrid clones for each. Mean ± SEM. One-way ANOVA. ns: p>0.05. **B.** Left: TRAP assay products for the indicated cell lines. Right: ddTRAP quantification of telomerase activity in the indicated cell lines. Copy number concentrations were normalized to the first replicate of 143B ρ^0^ cells (repl 1). Mean ± SEM. One-way ANOVA. **C.** qRT-PCR analysis of h*TR* and h*TERT* transcripts. Values were normalized first to *ACTB* mRNA levels and then to 143B ρ^0^ ratio. **D.** Left: Representative pictures of telomere dysfunction-induced foci (TIF) detection by co-localization of telomeres (red FISH probe) with 53BP1 DNA damage marker (green). White arrowheads indicate TIF. Blue: DAPI. Scale bar: 5 μm. Right: TIF quantification in clone #1 of all cybrids (first PDs post-selection) and in 143B ρ^0^ cells. n=47-63 nuclei. Mean ± SEM. Kruskal-Wallis test. ns: p>0.05. **E.** Left: Visualisation of telomere fusions (white arrowheads) in Cyb1 cells at PD7. 143B ρ^0^ and Cyb6 cells are shown as controls. Telomeres are detected by FISH (red) and DAPI stains DNA (blue). Scale bar: 5 μm. Right: Quantification of telomere fusions. The total number of analyzed chromosomes is shown for each cell line. χ^2^ test. ns: p>0.05.

Together, these findings demonstrate that mitochondrial DNA sequence variation exerts a strong influence on telomere length in 143B ρ^0^ recipient cells, ranging from telomere elongation to severe shortening.

### Low complex I activity shortens the telomeres of ρ^0^ recipient cells

We next investigated why different mtDNA sequences exerted such divergent effects on TL in 143B ρ^0^ recipient cells. When mitochondria are replenished in ρ^0^ cells during the formation of transmitochondrial cybrids, a shift from glycolysis to OXPHOS occurs (Fig S4A-B) and cells need to adapt to this metabolic change. Notably, the initial phase of ρ^0^ cell repopulation involves a temporary ROS burst, partly driven by incompletely assembled ETC supercomplexes that are known to elevate ROS levels (22). Complex I (CI) plays a crucial role in regulating the NAD^+^/NADH balance and this is crucial for the activity of NAD^+^-dependent poly (ADP-ribose) polymerase 1) (PARP1) repair enzyme, an enzyme required for the repair of telomeres undergoing oxidative stress in cultured cells (23–25). We thus hypothesized that optimal levels of CI activity may be required for telomere protection in cells experiencing acute oxidative stress during the *in vitro* metabolic shift before cellular adaptation. Consistent with this, we found significantly lower CI activity in Cyb1 and Cyb2 cybrids (Fig 3A) and a strong positive correlation between TL measured in the cybrids at the first passages and CI activity (Fig 3B). Supporting a lower CI activity in Cyb1 and Cyb2 cybrids, we noticed a significant up-regulation of the NAD^+^ salvage pathway genes *NAMPT* and *NAPRT1*, likely reflecting a compensatory mechanism (Fig 3C).

**Figure 3.**
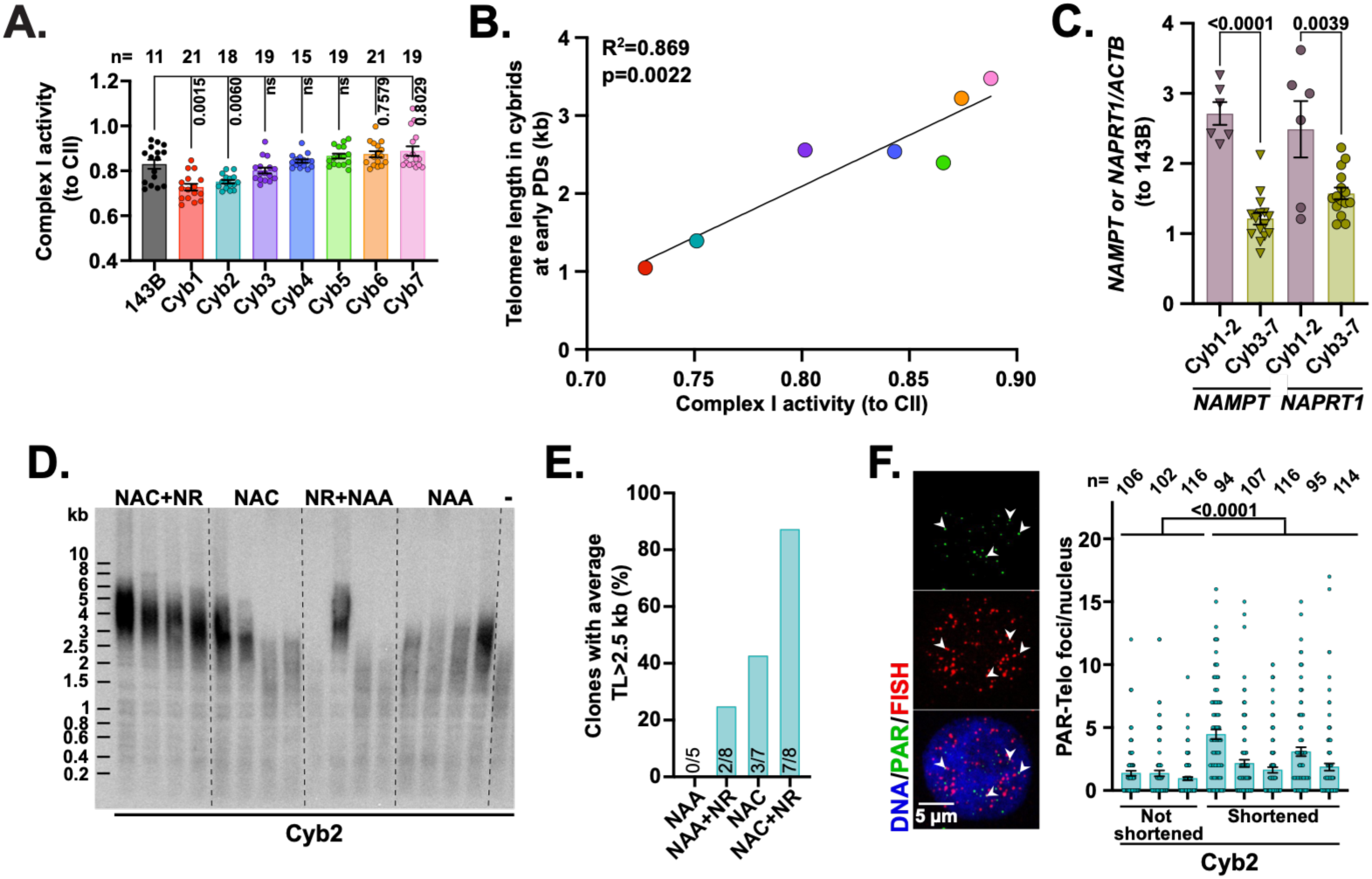
Telomere shortening in cybrids from donor with low complex I activity is counteracted by antioxidant and NAD^+^ precursor. **A.** Complex I (CI) activity in 143B and in the cybrids. Values were normalized to the activity of complex II (CII) which is entirely nuclear-encoded. n measurements from 2 biological replicates on clone #1 from all: at PD 4-7 or PD 16-24 post-selection. Mean ± SEM. Kruskal-Wallis test. ns: p>0.05. **B.** Correlation between TL in the cybrids at early PDs and average ratios of CI/CII activity. Color code as in A. **C.** Expression of *NAMPT* and *NAPRT1* genes from the NAD^+^ salvage pathway in cybrids (PD≥5 post-selection) from donors with either low (Cyb1 to 2) or normal (Cyb3 to 7) CI activity. Values were normalized to *ACTB* and to the ratios measured in 143B cells. Color code as in A. n=3 biological replicates. Mean ± SEM. Two-tailed Student’s *t* tests. **D.** Representative TRF analysis of TL in independent cybrid clones obtained by fusion between donor #2’s platelets and 143B ρ^0^ recipient cells in the presence of the indicated treatments or in the absence of added drug (-). 5 mM N-acetyl-L-cysteine (NAC), 5 mM N-acetyl-L-alanine (NAA, control) and/or 3 mM nicotinamide riboside (NR) were added at the time of the fusion and kept until clone isolation. **E.** Quantification of the TRF analyses conducted on independent cybrid clones isolated in the conditions detailed in D. The percentage of clones with average TL>2.5 kb was calculated. The specific number of independent cybrid clones analyzed for each condition is shown in the graph: NAA: 5, NAC: 7, NAC + NR: 8, NAA + NR: 8. **F.** Representative images and quantification of telomeric parylation in 8 independent Cyb2 clones obtained from 4 independent cybridisations, comparing cells that did or did not exhibit telomere shortening during the initial PDs following cybridisation (from panel D-E). Parylation was detected by IF (green) in the presence or PARG inhibitor; telomeres were detected by FISH (red) and DNA was stained with DAPI (blue). Scale bar: 5 μm. n=94-116 nuclei were counted for each clone. Mean ± SEM. Two-tailed Mann-Whitney test.

Building on the idea that, in the context of lower CI activity, telomeres may not be able to resist oxidative stress-induced shortening *in vitro*, we repeated the fusion of donor #2’s platelets to 143B ρ^0^ cells in the presence of N-acetyl-L-cysteine (NAC) antioxidant and/or nicotinamide riboside (NR), a precursor of NAD^+^. Strikingly, while telomeres were shortened in all clones (5/5) selected from cybrids obtained in the presence of N-acetyl-L-alanine (NAA) used as control, only one out of the eight selected cybrids obtained in the presence of both NAC and NR showed telomere shortening (Fig 3D-E). Treatment with either NAC or NR alone showed intermediate phenotype (Fig 3D-E). The detection of telomeric parylation signals further supported a role for NAD^+^-dependent PARP1 activity in repairing oxidative damage at shortened telomeres during the process of cybrid formation (Fig 3F).

Altogether, we propose that the *in vitro* process of cybrid formation exacerbates the impact of low CI activity on telomeres during the acute oxidative stress linked to a metabolic shift from glycolysis to OXPHOS.

### Lymphocyte TL inversely correlates with ROS levels in transmitochondrial cybrids

Having identified CI activity as a strong regulator of TL during *in vitro* cybridisation, we asked whether mtDNA-encoded CI activity could also account for the differences in TL observed in donors’ blood cells. However, analyzing age-adjusted lymphocyte TL against CI activity revealed no significant association (Fig 4A). This lack of correlation highlights the limitations of using the cybrid system to model the *in vivo* regulation of TL by CI activity.

**Figure 4.**
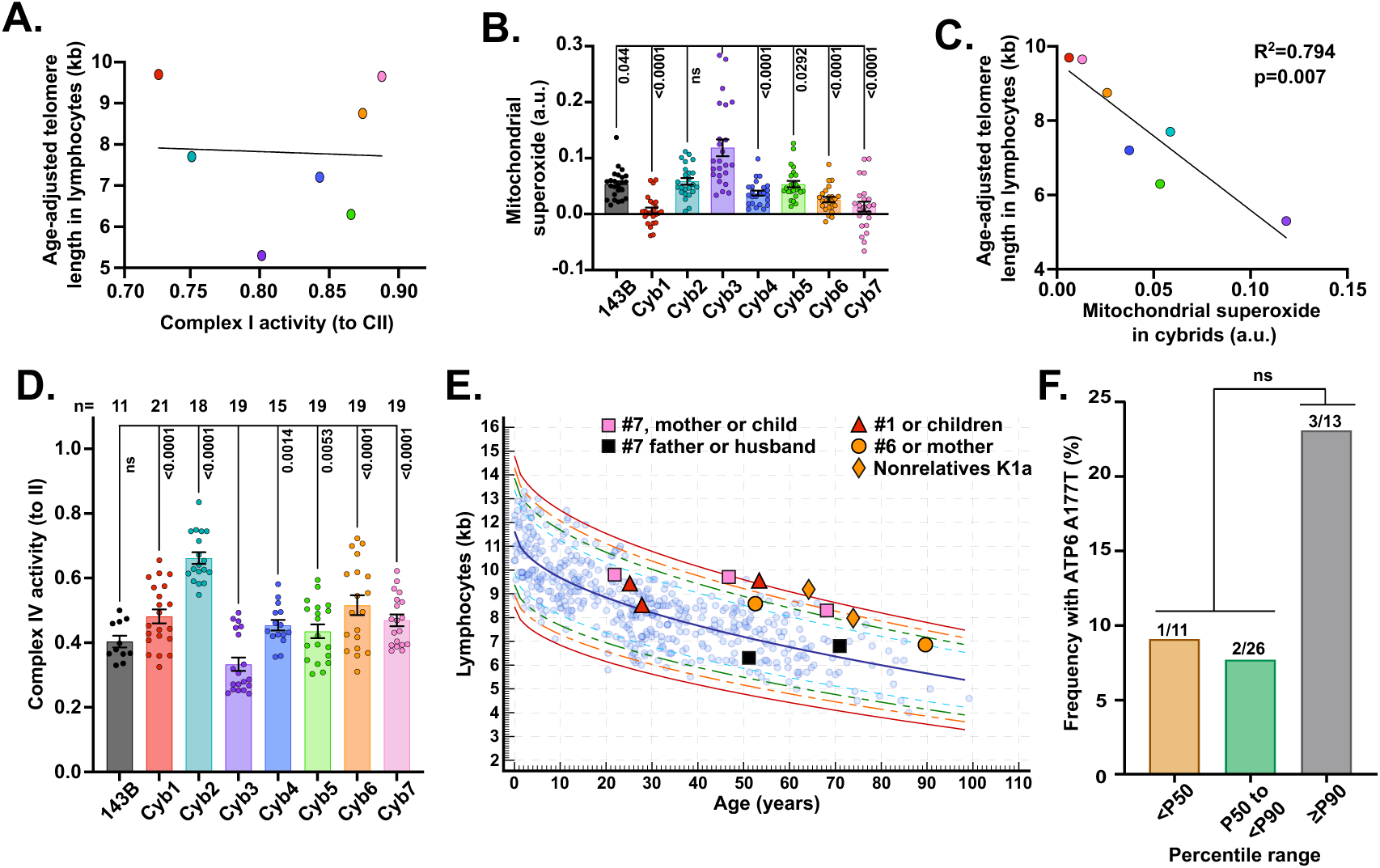
Mitochondrial superoxide production in cybrids inversely correlates with TL in lymphocytes. **A.** Lack of correlation between lymphocyte TL (in kb, adjusted to 50 yo) and average CI activity measured in cybrids from the seven corresponding donors and normalized to CII activity (CI/CII). TL in the donor lymphocytes was distributed as follows: >P90 (#1, #6 and #7), around P50 (#2 and #4), between P10 and P50 (#5) and at P10 (#3). **B.** Mitochondrial superoxide production measured by electron paramagnetic resonance and corrected for background using PEG-SOD2 in clone #1 from all cybrids. n measurements from 2 biological replicates with 3 technical replicates. Mean ± SEM. Kruskal-Wallis test. ns: p>0.05. **C.** Correlation between lymphocyte TL (in kb, adjusted to 50 yo) and average mitochondrial superoxide production measured in cybrids from the corresponding donors. **D.** Complex IV activity in 143B cells and in the cybrids. Values were normalized to CII activity. n measurements per cell line from 2 biological replicates. One-way ANOVA. ns: p>0.05. **E.** TL of donor #7 (H1b1e), donor #1 (T2b4a), donor #6 (K1a4a1) and their relatives is shown as indicated in the legend. Orange triangles show TL in unrelated K1a11b and K1a4a1a2b donors. **F.** Frequency of donors with the ATP6 A177T variant among n selected donors with lymphocyte TL in the indicated ranges (<P50, between P50 and <P90 and ≥P90). Chi-square test.

In mouse oocytes, impaired mitochondrial function, associated with elevated mitochondrial ROS production, adversely affects telomere elongation (10). In human cells, experimentally induced mitochondrial dysfunction was also reported to lead to telomere damage through the production of ROS (26). We therefore tested whether human mitochondrial genomes, through their impact on ROS levels, may modulate TL in blood cells. As transmitochondrial cybrids receive the donors’ mitochondria but no other genetic information, this cellular model allows to specifically assess the impact of the mitochondrial genome on ROS production. Mitochondrial superoxide production was measured by electron paramagnetic resonance (27) in clones #1 of all cybrids and in 143B cells (Fig 4B). We excluded 143B ρ^0^ cells from our superoxide measurements because these cells were grown in a different culture medium. Differences in ROS levels across the cybrids were not linked to distinct changes in oxidative *vs* glycolytic metabolism in the cybrids that were characterized by similar lactate/glucose and ATP/ADP ratios (Fig S5A-B). Strikingly, ROS levels in the cybrids inversely correlated with the age-adjusted TL measured by Flow-FISH in the corresponding donors’ lymphocytes (Fig 4C). Elevated levels of ROS were notably detected in Cyb3 cells. The mtDNA from donor #3 harbors the extremely rare F63L variant (m.T6090>C; Mitomap frequency: 0.002%) in the COX1 subunit, located within a transmembrane helix in close proximity to the heme A group and associated with reduced CIV efficiency (Fig 4D and Fig S6). Consistently, tracking TL over sequential population doublings revealed that Cyb3 cells are more vulnerable to progressive telomere shortening than Cyb6 or Cyb7 cells (Fig S7). Together, these findings support a role for mitochondrial ROS in human TL regulation.

### Rare mtDNA variants may account for the maternal inheritance of long telomeres

To further assess the relevance of our findings to a potential role of mitochondria in TL inheritance in humans, we searched for evidence of maternal inheritance of TL within families. Such analyses are most informative in families with extremely long (>P90) or short (<P10) telomeres. We therefore focused on three individuals with lymphocyte TL above the 90th percentile: donors #1, #6, and #7 (Fig 1C).

Despite donor #1 exhibiting lymphocyte TL at the 99^th^ percentile, TL in lymphocytes from both of this donor’s children was close to the 50^th^ percentile or only slightly above, arguing against a dominant maternal inheritance of the exceptionally long telomeres observed in donor #1 (Fig 4E). Donor #6 belongs to the K1a subhaplogroup Identified by the MitoAging project as one of the four mtDNA variants significantly associated with longevity, with the A177T variant also showing protective effect against Parkinson’s disease (28). Although we could not assess TL in donor #6’s progeny, as she does not have children, the fact that her mother showed telomeres at the 90^th^ percentile and that two European male donors, identified in independent cohorts (29) as having telomeres at, respectively, the 97.5^th^ and above the 99^th^ percentile (Fig 4E), were found to display K1a mtDNA, support the notion that this subhaplogroup is associated with long telomere inheritance. This led us to screen additional unrelated donors with varying TL for the presence of the m.G9055>A variant. The variant was identified in 1 of 11 donors with lymphocyte TL below the 50^th^ percentile, 2 of 26 donors with TL between the 50^th^ and 90^th^ percentiles, and 3 of 13 donors with TL at or above the 90^th^ percentile (Fig 4F and Fig S8), suggesting a trend toward higher frequency of the m.G9055>A (ATP6 A177T) variant among individuals with lymphocyte TL at or above the 90^th^ percentile (*p*=0.153). Further investigation will however be needed to correlate the ATP6 A177T with longer telomeres.

Donor #7 does not belong to the K1a haplogroup; nevertheless, both her mother and her daughter exhibited telomere lengths at the 95^th^ and near the 90^th^ percentile, respectively (Fig 4E). In contrast, the telomere lengths of her husband and her father were slightly above the 10^th^ and around the 50^th^ percentile, respectively (Fig 4E). These findings pointed to a maternal inheritance pattern associated with the H1b1e mtDNA subhaplogroup, identified through blood sequencing in these women but not previously linked to longevity. While the causal link between the rare m.A3796>G variant in donor #7 (ND1 T164A, CI) and the long telomere phenotype remains to be determined, this variant was absent in all 49 unrelated donors with varying telomere lengths that we screened. Collectively, these findings support the idea that multiple mitochondrial genome SNPs may contribute to the inheritance of telomere length.

## Discussion

While long telomeres are often viewed as markers of good cellular health, shorter leukocyte TL is linked to a higher risk of diseases affecting organs like the lungs and liver, as well as various cancers (4). Although aging is a major factor influencing TL, genetic variation also plays a key role and helps explain why TL established in the zygote strongly influences TL throughout life (7).

Unraveling the mechanisms that determine TL in the zygote is therefore highly relevant, particularly for optimizing *in vitro* fertilization strategies. A recent study in mice suggests that mitochondrial dysfunction impairs telomere elongation during embryonic development-a defect that was largely reversed by antioxidant treatment, highlighting the harmful effect of ROS on TL regulation in the embryo (10). In humans, prenatal exposure to psychological stress has also been linked to shorter leukocyte TL in offspring (30). Given the established connection between psychological and oxidative stress (31,32), these findings point to a similar influence of mitochondrial function on TL regulation during human embryogenesis. Building on these findings -and previous evidence suggesting stronger maternal inheritance of TL in humans (8)-we hypothesized that the mitochondrial genome may influence human TL. To investigate this, we employed a transmitochondrial cybrid model using platelets from seven donors with known leukocyte TL and distinct mitochondrial subhaplogroups. Notably, we observed an inverse correlation between donor leukocyte TL and mitochondrial ROS levels in the cybrids, indicating that mitochondrial subhaplogroup affects mitochondrial function and suggesting a potential role for mtDNA variants, such as ATP6 A177T, in ensuring longer leukocyte TL *in vivo*. Strikingly, 32.7% of the genetically homogeneous population of Ashkenazi Jews express the ATP6 A177T variant (12). As telomeres from Ashkenazy centenarians blood cells are longer than in control groups (33), it will be interesting to test the link to the m.G9055>A variant encoding ATP6 A177T. While the cybrid system has limitations in modeling *in vivo* TL regulation by CI activity, our findings reveal a notable contrast among donors. Donors #6 and #7, who show signs of maternal inheritance for their long telomeres, exhibit high mitochondrial CI activity. In contrast, donor #1, who also has long telomeres but low mtDNA-encoded CI activity, did not transmit her long telomeres to her offspring. In summary, our data indicate that low levels of mitochondrial ROS, combined with potent CI activity, may be key for a mtDNA-driven maternal inheritance of long telomeres. The Japanese m.C5178A longevity-associated variant in the *ND2* gene (34) further supports an important role for CI in promoting mitochondrial fitness, and possibly TL regulation. Larger studies will however be needed to support the link between mtDNA genome and leukocyte TL.

Whether and how the centenarian-linked K1a mtDNA contributes to reduced ROS levels awaits further investigation. As a defective aerobic production of ATP, linked to the ATP6 A177T variant, was proposed to account for the lack of mtDNA K haplogroup in Finnish elite endurance athletes (35), we hypothesize that, by reducing ATP aerobic production, ROS production decreases, thus providing a possible explanation to this longevity-associated mtDNA (36). Consistent with this, Cyb6 cells showed a distinct response to FCCP uncoupling agent as, unexpectedly, respiration could not be restored in cells with oligomycin-inhibited ATP synthase, suggesting a defective ability in increasing oxygen consumption rate to face an energy demand increase (Fig S4A-B). This could be related to the location of the A177T variant in a crucial region of ATP6, along the proton pore (37).

Experimentally induced mitochondrial dysfunction leads to telomere damage (26). Mechanistically, the *TTAGGG* repeats of telomeres are exquisitely sensitive to 8-oxo-guanine formation, a lesion recognized by the base excision repair pathway that, after removal of the oxidized base, generates single-strand breaks eventually recruiting NAD+-dependent PARP1 repair enzyme (24,38). If left unrepaired in replicating cancer cells, these breaks can convert into double-strand breaks, resulting in distal telomere loss and chromosomal fusions (25). In normal fibroblasts and epithelial cells, telomeric 8-oxoG lesions induced by a chemoptogenetic tool were sufficient to activate p53-dependent senescence (39), highlighting the harmful effects of oxidative stress at telomeres on cellular function. During the generation of transmitochondrial cybrids, cells undergo a metabolic shift from glycolysis to OXPHOS, accompanied by increased ROS production. Before cells adapt to this shift, oxidative stress can be particularly damaging. Mitochondrial platelets with low CI activity triggered significant telomere shortening in recipient cells during early cybrid formation -a loss that was prevented by co-treatment with an antioxidant and an NAD⁺ precursor. Although we did not determine whether the rescue was dependent on PARP1, these results support the essential role of CI in maintaining PARP1 activity *via* regulation of the NAD⁺/NADH balance. Although the observed telomere loss may be exacerbated by the *in vitro* oxidative stress conditions-since donors with low CI activity did not show short leukocyte telomeres-our findings are consistent with reports that NAD⁺ supplementation can alleviate telomere dysfunction in cells from dyskeratosis congenita patients (40) and improve hematopoietic function and telomere integrity in Tert⁻/⁻ mice (41). Data are also consistent with the observation that mice lacking PARP display telomere shortening and chromosomal instability (42). Together, these data bring support to the hypothesis that, when extremely pathogenic and associated with high oxidative stress levels, such as in MELAS or LHON patients (43), mutations in CI may impair telomere length resetting *in utero*.

Our study also unexpectedly revealed a reduction in telomere length in 143B cells following mitochondrial depletion. This telomere shortening was associated with decreased telomerase activity, as measured by ddTRAP, but not with reduced expression of the telomerase subunits h*TERT* or h*TR*. 143B osteosarcoma cells are deficient in thymidine kinase 1 and therefore lack an effective pyrimidine salvage pathway. In the absence of oxidative phosphorylation, the *de novo* thymidine synthesis pathway is also compromised, rendering 143B ρ^0^ cells dependent on uridine supplementation for survival. Although this hypothesis requires further investigation, we propose that the shorter telomeres observed in 143B ρ^0^ cells may be linked to the emerging role of thymidine nucleotide metabolism in the regulation of human telomere length, potentially through an allosteric effect of dTTP on telomerase activity (44–46).

Altogether, our findings that certain mitochondrial genomes may support telomere elongation are relevant to three-parent *in vitro* fertilization, a technique that uses donor eggs to prevent the transmission of mitochondrial diseases from mother to child (47). Future work will be necessary to investigate how mitochondrial activity may be involved and whether the nucleotide metabolism that recently emerged as an important regulator of telomerase activity may play additional role.

## Materials and Methods

### Human study participants

Blood samples were obtained from a total of 491 healthy donors. Cohort characteristics are as follows: 261 females and 230 males, median age of 26.3 y (7 days-99.2 y). Protocols were approved by the CHEF institutional Ethics committee and written informed consents were obtained from all 18 year-old or over volunteers. For children below 18, left-over blood samples were obtained through the Cliniques universitaires Saint-Luc after screening for inclusion.

### Leukocyte purification

Blood samples were collected in EDTA tubes and kept at room temperature for a maximum of 48 h before processing. Leukocytes were purified after erythrocyte lysis with ice-cold 0.155 M NH_4_Cl (Sigma-Aldrich), 0.01 M KHCO_3_ (Sigma-Aldrich) and 0.1 mM EDTA (Sigma-Aldrich) solution as described (14). Leukocytes were then resuspended in hybridization buffer (5 % dextrose/10 mM HEPES/0.1 % BSA), and frozen at −80 °C in 80 % FBS (Cytiva)/20 % DMSO (Sigma-Aldrich).

### Flow-FISH

The average telomere length (TL) was measured in peripheral blood leucocytes by flow cytometry combined with FISH (Flow-FISH), using bovine thymocytes as reference as previously described (14). TL of bovine thymocytes was measured by telomere restriction fragment (TRF) analysis (see below). TL was evaluated in lymphocyte and granulocyte populations distinguished by LDS751 staining of DNA as described (14). The telomeric PNA probe was as follows: FITC-OO-CCCTAACCCTAACCCTAA (Panagene). Values obtained for TL in lymphocytes and granulocytes are provided in Table S1.

### Genomic DNA extraction

Genomic DNA was extracted by overnight digestion at 45°C with 100 µg/mL proteinase K (Sigma-Aldrich) in lysis buffer (10 mM Tris–HCl, 10 mM EDTA, 1 % SDS; pH 8.0) followed by DNA purification with 25:24:1 phenol–chloroform–isoamyl alcohol (Sigma-Aldrich) and precipitation in isopropanol (Thermo Fisher Scientific) and 0.3 M sodium acetate; pH 5.5 (Sigma-Aldrich). gDNA was subsequently treated with 0.2 mg/mL RNAse A (Thermo Fisher Scientific) for 1 h at 37 °C, purified, and precipitated again as described above. DNA concentration, purity and integrity were evaluated with Nanodrop (Thermo Fisher Scientific).

### MtDNA subhaplogroup identification

PCR reactions were performed on 20-50 ng total DNA using GoTaq® G2 DNA Polymerase (Promega) and the primers provided in Table S2 so as to cover the entire mtDNA sequence (47). PCR program was as follows: 3 min at 95 °C followed by 40 cycles of 95 °C for 45 s, 47/50/55 °C for 45 s, and 72 °C for 1 min, with a final step at 72 °C for 5 min. PCR products were loaded on 1 % agarose gel (Eurogentec), stained with 0.5 µg/mL ethidium bromide (Sigma-Aldrich) and extracted from the gel using SmartPure Gel Kit (Eurogentec), according to the manufacturer’s instructions, followed by Sanger sequencing (Azenta/Genewiz). MtDNA subhaplogroups were identified using MITOMASTER tool on mitomap.org website.

### MtDNA content quantification

MtDNA content quantification was performed by qPCR on 60 ng of total DNA, using KAPA SYBR FAST (Sigma-Aldrich) and primer sequences (48) provided in Table S2.

### Cell culture and treatments

143B cells and cybrids were grown in DMEM high glucose (4.5 g/L) supplemented with 10 % FBS (Cytiva) and 1 % Penicillin/Streptomycin (Capricorn Scientific). 143B ρ^0^ cells were grown in DMEM high glucose supplemented with 0.11 g/L pyruvate (Thermo Fisher Scientific), 10 % FBS (Cytiva), 50 μg/mL uridine (Sigma-Aldrich) and 1 % Penicillin/Streptomycin (Capricorn Scientific). For the subcloning of 143B ρ^0^ population, cells were treated exactly as if they would undergo fusion with platelets and kept throughout the cloning process in ρ^0^culture medium. All cell lines were regularly checked for mycoplasma contamination. For this, 100 µL of culture medium were heated at 95°C for 5 min before spinning down. qPCR reactions were performed on 2 µL of the supernatant as described previously (49) using primers listed in Table S2.

### Isolation of human platelets

Blood was collected in 8.5 mL-CPDA tubes (Sarstedt) and centrifuged at 330 x *g* for 20 min at 22 °C. The upper platelet-rich plasma fraction was treated with 4 µg/mL eptifibatide (GlaxoSmithKline) and 1 U/mL apyrase (Sigma-Aldrich) on ice before centrifugation at 800 x *g* for 10 min at room temperature. The pellet was resuspended into 5 mL physiological saline solution containing 4 µg/mL eptifibatide and 1 U/mL apyrase. Platelets were counted and diluted with the saline solution to obtain 10^5^ platelets/µL and stored for 30-45 min at 37 °C. 40 x 10^6^ platelets were then transferred into a 2 mL Eppendorf tube, centrifuged at 800 x *g* for 10 min at room temperature and directly processed for fusion with 143B ρ^0^ cells.

### Cybrid formation

Cybrids were obtained as previously described (21). A detailed protocol is provided as Supplementary information.

### Generation of new 143B ρ^0^ clones

New 143B ρ^0^ clones were generated as previously described (50). Briefly, 1 × 10⁵ cells were seeded in 10-cm dishes and allowed to adhere overnight at 37 °C. Cells were then cultured in DMEM, High Glucose, Pyruvate (Gibco, 41966029) supplemented with 10% FBS, 1% penicillin–streptomycin, 50 µg/ml freshly added uridine (Sigma, U3003), and 50 ng/ml ethidium bromide (Invitrogen, 15585011). The treatment was maintained for 4 weeks prior to clonal isolation. MtDNA depletion in the resulting clones was checked by qPCR as described above.

### Mitochondrial superoxide assessment by EPR spectroscopy

The electron paramagnetic resonance (EPR) assay is based on the oxidation of Mito-TEMPO-H, a diamagnetic cyclic hydroxylamine that accumulates inside the mitochondria and is transformed into a paramagnetic nitroxide. To isolate the contribution of superoxide to the oxidation of the probe, PEG-SOD2 was used as a control as described previously (27,51–53). A detailed protocol is provided as Supplementary information.

### Seahorse analyses

Oxygen consumption rates (OCR) were measured using a Seahorse XFe96 plate reader (Agilent). All assays were carried out using a seeding density of 10,000 cells/well in non-buffered DMEM, adjusted at pH 7.4. Mitochondrial function parameters (*i.e.* basal and maximal respirations and ATP production-linked OCR) were evaluated, with the XF Cell MitoStress Test (Agilent), after sequential treatment with 1 µM oligomycin, 1 µM FCCP and 0.5 µM rotenone/antimycin A, as previously described (54). ETC complex activity was assessed as previously reported (55), with slight modifications. Briefly, 10^4^ cells per well were first permeabilized with 1 nM Seahorse XF Plasma Membrane Permeabilizer (Agilent) and respiration was stimulated by adding 10 mM pyruvate, 5 mM malate, and 2 mM ADP. After the baseline scan, 100 nM rotenone was then added to inhibit complex I and 10 mM succinate was injected to establish complex II respiration. Last, 4 μM antimycin A was added to inhibit complex III and 100 μM N,N,N’,N’-Tetramethyl-p-phenylenediamine dihydrochloride (TMPD) + 10 mM ascorbate was injected as a complex IV substrate. OCR data were then normalized according to protein content in each experimental well.

### Telomere restriction fragment (TRF) length analysis

TRF analyses were performed as described previously (56) on 2-5 µg of RNAse-treated genomic DNA digested with 20 U *Hinf*I and 20 U *Rsa*I and using [γ-^32^P] ATP-labelled (TAACCC)_4_ probe. Smart Ladder (Eurogentec) was run together with the samples, stained with ethidium bromide and imaged with a ruler. TRF were quantified using the online available WALTER (Web-based Analyser of the Length of Telomeres) software (https://www.ceitec.eu/chromatin-molecular-complexes/rg51/tab?tabId=125#WALTER).

### TeSLA analysis

The procedure described previously (57) was followed for TeSLA analysis. Primer sequences are provided in Table S2.

### Immunofluorescence and fluorescence *in situ* hybridization

Immunofluorescence and FISH experiments were performed as described previously (56) using the following antibodies and telomeric probe: rabbit anti-53BP1 (Novus Biologicals), Alexa 488 goat anti-rabbit (Invitrogen), mouse anti-Poly(ADP-ribose) (Millipore), Alexa 488 Goat anti-mouse (Abcam), 5’(TYE 563)GGGTtAGGGttAGgGTTAGGGttAGGGttAGGGtTA(TYE 563) (small letters indicate LNA^TM^ modified bases) (Qiagen). For the detection of parylated proteins, cells were grown in the presence of 1 μM PDD00017273 PARG inhibitor (Sigma-Aldrich). For telomeric FISH on metaphase spreads, cells were treated as described previously (58) and telomeres were probed with 5’(TYE 563)GGGTtAGGGttAGgGTTAGGGttAGGGttAGGGtTA(TYE 563). Images were acquired with the Cell Observer Spinning Disk confocal microscope (Zeiss) with 100× objective and analyzed using ImageJ software (National Institute of Health), while maintaining the same threshold for samples from the same experiment.

### Liquid chromatography-mass spectrometry analysis of metabolites

6-well plates with cultured cells were rapidly washed twice with ice cold water and submerged in liquid nitrogen. 300 µL of a solution of 90 % methanol and 10 % chloroform were added, lysates were scraped and transferred to microcentrifuge tubes for centrifugation for 15 min at 4 °C and 22, 000 x *g*. The supernatants were dried in a Speed-Vac vacuum concentrating system and resuspended into 50 µL of 50:50 water:methanol solution. Analyses by LC-MS were performed as previously described (59) based on a method by Coulier *et al.* (60). A detailed protocol is provided as Supplementary information.

### Dosage of extracellular glucose and lactate

Glucose and lactate concentrations were measured in extracellular media by using specific enzymatic assays and an ISCUSflex microdialysis analyzer (M Dialysis), as previously described (54).

### RNA isolation and reverse transcription quantitative real-time PCR

RNA was extracted using TriPure Isolation Reagent (Sigma-Aldrich) according to the manufacturer’s instructions. For cDNA synthesis, 1 μg of RNA was retro-transcribed with 200 U MMLV-RT (Thermo Fisher Scientific), 0.2 µg random hexamers (Thermo Fisher Scientific), 20 U RiboLock RNase inhibitor (Thermo Fisher Scientific) and 2 mM dNTP mix. qPCR reactions were performed using KAPA SYBR FAST (Sigma-Aldrich) and the following program: 95°C for 10 min followed by 40 cycles of 95 °C for 15 sec and 60 °C for 30 sec. Primer sequences are provided in Table S2.

### Telomerase repeat amplification protocol (TRAP) and droplet digital TRAP (ddTRAP)

For telomerase activity assessment *in vitro* using the TRAP assay, 500,000 cells were lysed in 100 µL CHAPS lysis buffer (TRAPeze S7700, Millipore) on ice for 30 min and proteins were quantified by Bradford assay (Bio-Rad). 500 ng of the lysis mixture were subjected to the TRAP assay following the Millipore kit instructions and TRAP products were separated on TBE/acrylamide:bisacrylamide (19:1) gel and visualized by staining with SYBR Gold Nucleic Acid GelStain (Invitrogen) with a PhosphoImager (Fujitsu). For ddTRAP (61), 500 ng of the lysis mixture were added to 50 µL of 1× TRAP buffer (200 mM Tris–HCl, pH 8.3, 15 mM MgCl_2_, 2.5 mM each dNTPs, 200 mM TS primer (Table S2), and 0.4 mg/ml BSA), and extension reaction was performed by incubating the mix at 25°C for 40 min, followed by deactivation at 95°C for 5 min. For the ddPCR reaction, 3.75 μL of lysis-extension mix were added to 1x EvaGreen QX200 Super-mix (Bio-Rad), 50 nM TS primer, 50 nM ACX primer (Table S2) and H_2_0 up to 25 µL. PCR reaction was performed as follows: initial denaturation for 5 min at 95°C followed by 40 cycles of 95°C/30 s, 54°C/30 s, and 72°C/30 s. Data were analyzed on Quantasoft software.

### Statistics

Statistical analyses were performed using PRISM 8 software. Normality of the data was always checked before statistical tests. Details are provided in the Figure legends. Nomograms for Flow-FISH-based measurements of TL in the healthy population were drawn using the Age-related reference interval function of MedCalc Software (Belgium) based on methods described previously (62–64).

## Supporting information

All SI

## Acknowledgments

We are grateful to all volunteers for providing blood samples. We thank Sandrine Horman, Marie Octave and Valentine Robaux for help with platelet purification and Géraldine Aubert and Mary Armanios for guidance in Flow-FISH. We are grateful to Victoria Van Regemorter and Caroline Huart for sharing unpublished data on TL in their cohort. We thank Chloé Despontin and Antoine Fattaccioli for technical support, Céline Coquette for help with blood collection and Mélina Vaurs, and Nausica Arnoult for critical reading of the manuscript. This work was supported by the Fonds National pour la Recherche Scientifique (FNRS) (Belgium), The Belgian Foundation against Cancer and WELBIO (Walloon Excellence in Life Sciences and Biotechnology) (grant X250624 to CC). Patrick Revy’s team is Labellisé Ligue (France). We are grateful to the de Duve Institute for constant support.

