## Supplementary material for "Human mitochondrial DNA variants influence telomere length: evidence from a transmitochondrial cybrid model": All SI

**This PDF file includes:**

Supporting text
Figures S1 to S8
SI Reference
Tables S1 to S2

**Supporting Information Text**

### Supplementary methods

#### Cybrid formation

For cybrid formation, $10^6$ exponentially growing 143B $\rho^0$ cells (Prof. G. Attardi, California Institute of Technology, USA) were trypsinized and centrifuged at 290 x *g* for 5 min before resuspension in 2 mL calcium-free DMEM (Thermo Fisher Scientific). 143B $\rho^0$ cells were slowly added, without mixing, to the freshly-isolated platelet pellet before centrifugation at 200 x *g* for 10 min at RT. Supernatant was discarded and pellet was resuspended in 100 µL PEG 1500 (Sigma-Aldrich) fusion solution (490 µL calcium-free DMEM, 10 µL DMSO, 0.5 g PEG 1500). After homogenization during exactly 1 min, 10 mL DMEM high glucose/pyruvate supplemented with 10 % FBS, 50 µg/mL uridine, 100 µg/mL BrdU (Sigma-Aldrich) and 1 % Penicillin/Streptomycin were added and the cellular suspension was directly distributed into 5 Petri dishes containing 8 mL of the same medium. After 3-5 days, medium was changed and cybrids were selected in DMEM high glucose supplemented with 10 % dialyzed FBS (Thermo Fisher Scientific), 100 µg/mL BrdU and 1 % Penicillin/Streptomycin for 10 days. Colonies from the 5 Petri dishes were isolated with cloning cylinders and transferred into 24-well plates containing DMEM high glucose supplemented with 10 % FBS and 1 % Penicillin/Streptomycin for expansion. Clones were then transferred into flasks and counted for the first time when they had reached 1 to 10 million cells. This time point initiated the calculation of population doublings (PDs), although we estimate that the clones had already gone through at least 19-23 PDs post-fusion. The same procedure was followed for the second fusion of 143B $\rho^0$ cells with freshly-isolated platelets from donor #2. Five mM N-Acetyl-L-alanine (NAA, Sigma-Aldrich) or N-Acetyl-L-cysteine (NAC, Sigma-Aldrich) and/or 3 mM Nicotinamide riboside (NR, Sigma-Aldrich) were added directly after the 1 min incubation with PEG. Medium was replaced every 3 days and drug treatments were maintained until the transfer of the isolated colonies into T25 flasks.

#### Mitochondrial superoxide assessment by EPR spectroscopy

We used a Bruker EMX-Plus spectrometer operating in X-Band (9.85 GHz) and equipped with a PremiumX ultra-low noise bridge and a SHQ high sensitivity resonator. During all the experiments, the EPR cavity was kept at 310 K with a continuous air flow (400 L/h). A mixture was prepared by mixing 37 µL of cell suspension (previously harvested, stock solution $1.5x10^7$/mL of the appropriate medium), 0.5 µL of diethylenetriaminepentaacetic acid (DTPA) (100 mM), 5 µL of PBS (pH 7.4) and 7.5 µL of MitoTempoH (1 mM). The Mito-TEMPO-H (1-hydroxy-4-[2-triphenylphosphosphonio)-acetamido]-2,2,6,6- tetramethylpiperidine) solution (Enzo Life Sciences) was flushed with argon before each experiment to avoid probe oxidation. Control measurements were performed after incubation for 15 min with 200 U/mL PEG-SOD2 (Sigma-Aldrich). The final mixture was transferred in a 12-cm long gas-permeable polytetrafluoroethylene (PTFE) tubing (Zeus) (inside diameter 0.025 in, wall thickness 0.002 in) and folded to be inserted into an open quartz tube. The EPR parameters set in Bruker Xenon Spin fit program were: microwave power, 20 mW; modulation frequency, 100 kHz; modulation amplitude, 0.1 mT; center field, 336.5 mT; sweep width, 1.5 mT; sweep time, 30.48 s. The first EPR acquisition was performed 3 min after the probe was added to the cell mixture. For each biological replicate, measurements were done in triplicates. Measurements were performed on cybrids (clone #1 for all) between PD43 and PD63. Data were analyzed by performing a double integration on selected regions of peaks and retrieving background values obtained with PEG-SOD2. Full details of the procedure are published elsewhere (1).

**Liquid chromatography-mass spectrometry analysis of metabolites**

Five µL of the 50 µL 50:50 water:methanol solution were analyzed with an Inertsil 3 µm ODS-4 column (150 x 2.1 mm; GL Biosciences) at a flow rate of 0.2 mL/min using an Agilent 1290 HPLC system coupled to an Agilent 6550 Q-TOF MS in negative mode. Mobile phase A consisted of 5 mM hexylamine adjusted to pH 6.3 with acetic acid and phase B of 90 % methanol/10 % 10 mM ammonium acetate adjusted to pH 8.5 with ammonia. The mobile phase profile was set up as follows: 0 – 2 min at 0 % B; 2 – 6 min from 0 to 20 % B; 6 – 17 min from 20 to 31 %B; 17 – 36 min from 31 to 60 % B; 36 – 41 min from 60 to 100 % B; 41 – 51 min at 100 % B; 51 – 53 min from 100 to 0 % B; 53 – 60 min at 0 % B. Compound identification was based on exact mass (<5 ppm) and retention time compared to standards. The areas under the curve of extracted-ion chromatograms of the $[M\text{-}H]^-$ forms were determined using MassHunter software (Agilent) and normalized to the mean value of a batch of 150 other metabolites (‘total ion current’).

**Supplementary figures**

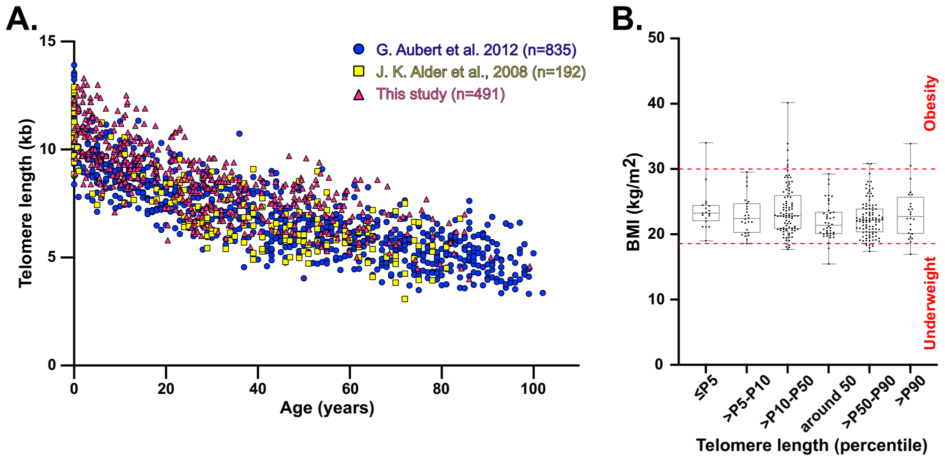

A.
G. Aubert et al. 2012 (n=835)
J. K. Alder et al., 2008 (n=192)
This study (n=491)
Telomere length (kb)
15
10
5
0
0
20
40
60
80
100
Age (years)
B.
BMI (kg/m2)
50
40
30
20
10
0
Obesity
Underweight
≤P5
>P5-P10
>P10-P50
around 50
>P50-P90
>P90
Telomere length (percentile)

**Fig. S1. Analysis of Flow-FISH reference curves. A.** Comparison of Flow-FISH curves obtained in lymphocytes from healthy donors by Alder *et al.* (black squares), Aubert *et al.* (red circles) and us (blue triangles). **B.** Body mass index (BMI) was calculated in donors classified according to the percentile of TL in their lymphocytes as indicated. Min, Lower quartile, Median, Upper quartile, Max. None of the comparisons gave significant difference using the Kruskal-Wallis test. ns: $p>0.05$.

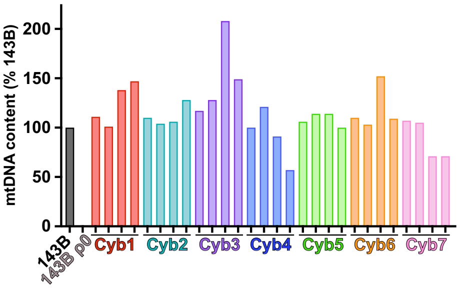
mtDNA content (% 143B)
200
150
100
50
0
143B
143B ρ0
Cyb1
Cyb2
Cyb3
Cyb4
Cyb5
Cyb6
Cyb7

**Fig. S2. mtDNA quantification in the cybrids.** Quantification of mtDNA content by qPCR in four independent cybrid clones from the seven donors at early PDs, normalized to h*TR* genomic DNA content and to 143B.

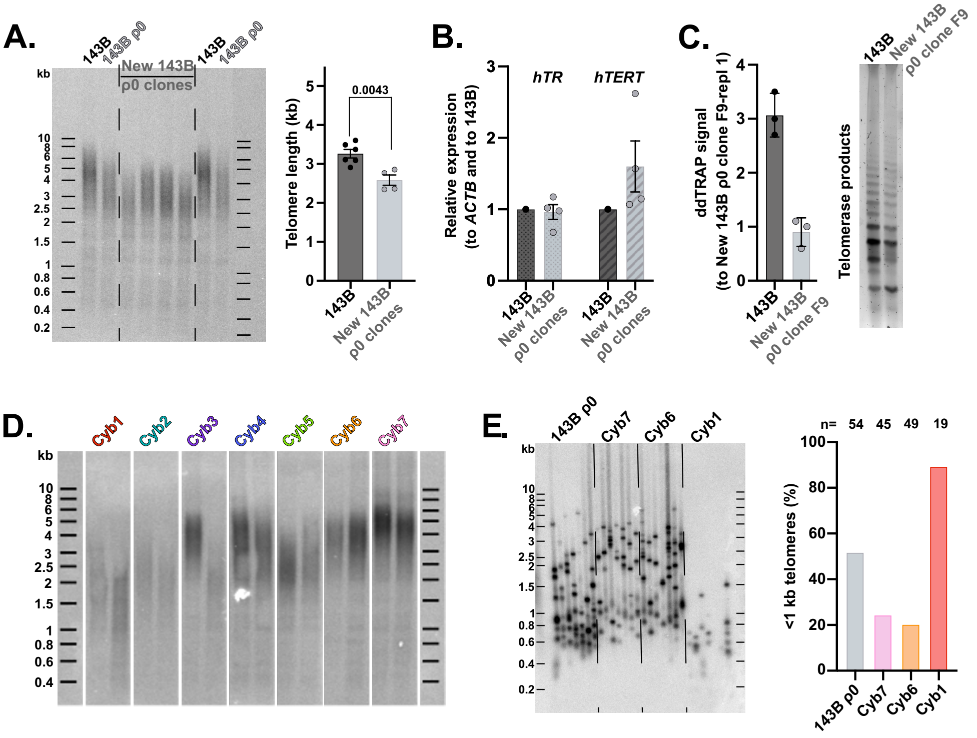
A.
143B
143B ρ0
New 143B ρ0 clones
143B
143B ρ0
kb
Telomere length (kb)
0.0043
143B
New 143B ρ0 clones
B.
Relative expression (to ACTB and to 143B)
hTR
hTERT
143B
New 143B ρ0 clones
C.
ddTRAP signal (to New 143B ρ0 clone F9-repl 1)
143B
New 143B ρ0 clone F9
Telomerase products
D.
Cyb1
Cyb2
Cyb3
Cyb4
Cyb5
Cyb6
Cyb7
E.
143B ρ0
Cyb7
Cyb6
Cyb1
n= 54 45 49 19
<1 kb telomeres (%)
143B ρ0
Cyb7
Cyb6
Cyb1

**Fig. S3. Mitochondria depletion in 143B cells shortens telomeres and reduces telomerase activity. A.** Left: Representative Southern blot (TRF) analysis of telomeres in 143B, 143B $\rho^0$ (Prof. G. Attardi, California Institute of Technology, USA) and four clones from the newly generated 143B $\rho^0$ cells. Right: Quantification of A. Additional TL values for 143B cells were included. Mean ± SEM. Unpaired Student's *t* test. **B.** qRT-PCR analysis of h*TR* and h*TERT* transcripts in the four new 143B $\rho^0$ clones shown in panel A. Values were normalized first to *ACTB* mRNA levels and then to 143B ratio. Mean ± SEM. **C.** Left: ddTRAP quantification of telomerase activity in 143B and new 143B $\rho^0$ clone F9 cell extracts. Copy number concentrations were normalized to the first replicate of new 143B $\rho^0$ clone F9 (repl 1). Mean ± SEM. Right: TRAP assay products for the same cell lines. **D.** Representative TRF analysis of telomeres in two independent cybrid clones from each donor at the first PDs post-selection (related to Fig 2A). **E.** Left: TeSLA analysis of TL in clone #1 of the indicated cybrids at similar PD values and in 143B $\rho^0$ recipient cells. n=1. Right: Frequency of telomeres with length <1kb in 143B $\rho^0$ cells, Cyb7, Cyb6 and Cyb1 cybrids calculated on n telomeres.

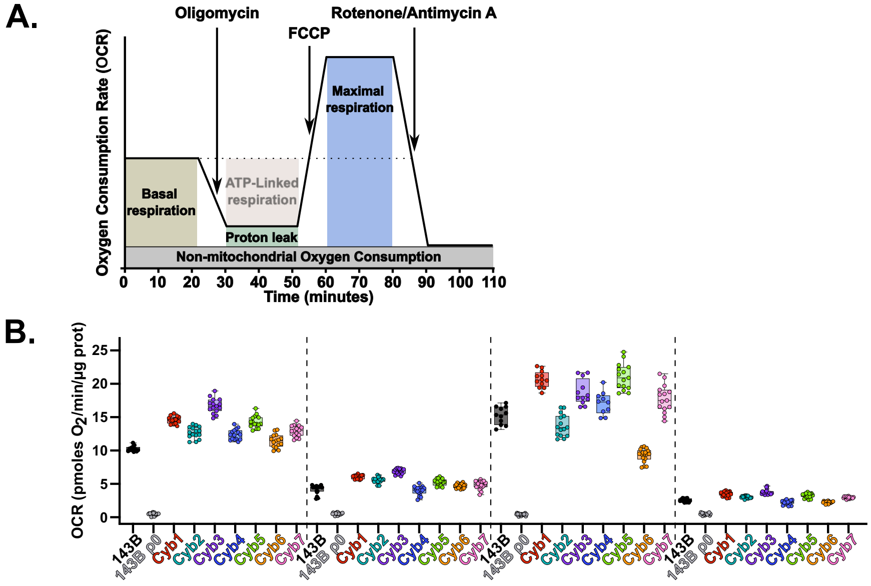
A.
Oligomycin
FCCP
Rotenone/Antimycin A
Maximal respiration
Basal respiration
ATP-Linked respiration
Proton leak
Non-mitochondrial Oxygen Consumption
Oxygen Consumption Rate (OCR)
Time (minutes)
B.
OCR (pmoles O2/min/µg prot)
143B
143B ρ0
Cyb1
Cyb2
Cyb3
Cyb4
Cyb5
Cyb6
Cyb7

**Figure S4. Oxygen consumption rate (OCR) measurement in cybrids. A**. Overview of the Seahorse Cell Mito Stress Test used to evaluate basal and maximal OCR, the ATP-linked OCR and the proton leak contribution to OCR. **B.** OCR values in 2-4 different cybrid clones from each donor in the conditions described in A. n=2-5 biological replicates with technical replicates. Mean ± SEM.

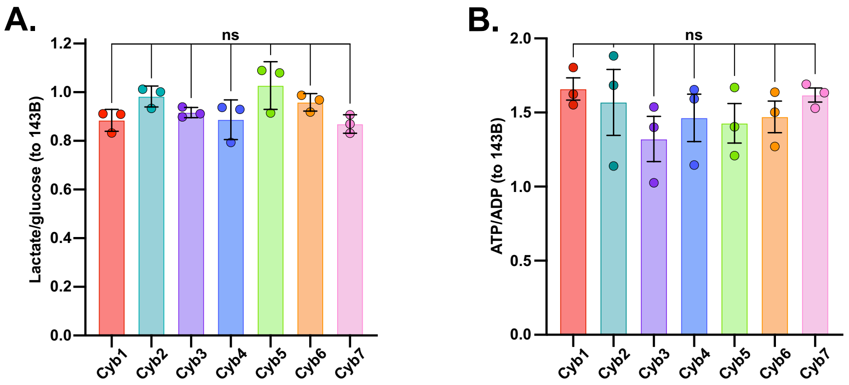
A.
ns
Lactate/glucose (to 143B)
1.2
1.0
0.8
0.6
0.4
0.2
0.0
Cyb1
Cyb2
Cyb3
Cyb4
Cyb5
Cyb6
Cyb7
B.
ns
ATP/ADP (to 143B)
2.0
1.5
1.0
0.5
0.0
Cyb1
Cyb2
Cyb3
Cyb4
Cyb5
Cyb6
Cyb7

**Figure S5. Similar metabolic activity of the cybrids. A.** Lactate and glucose concentrations measured by enzymatic colorimetric assay in clones #1 of the cybrids. Lactate/glucose ratios were normalized to 143B. n=3 biological replicates. Mean ± SEM. **B.** ATP and ADP levels measured by LC/MS in clones #1 of cybrids. n=3 biological replicates. ATP/ADP ratios were normalized to 143B. Mean ± SEM.

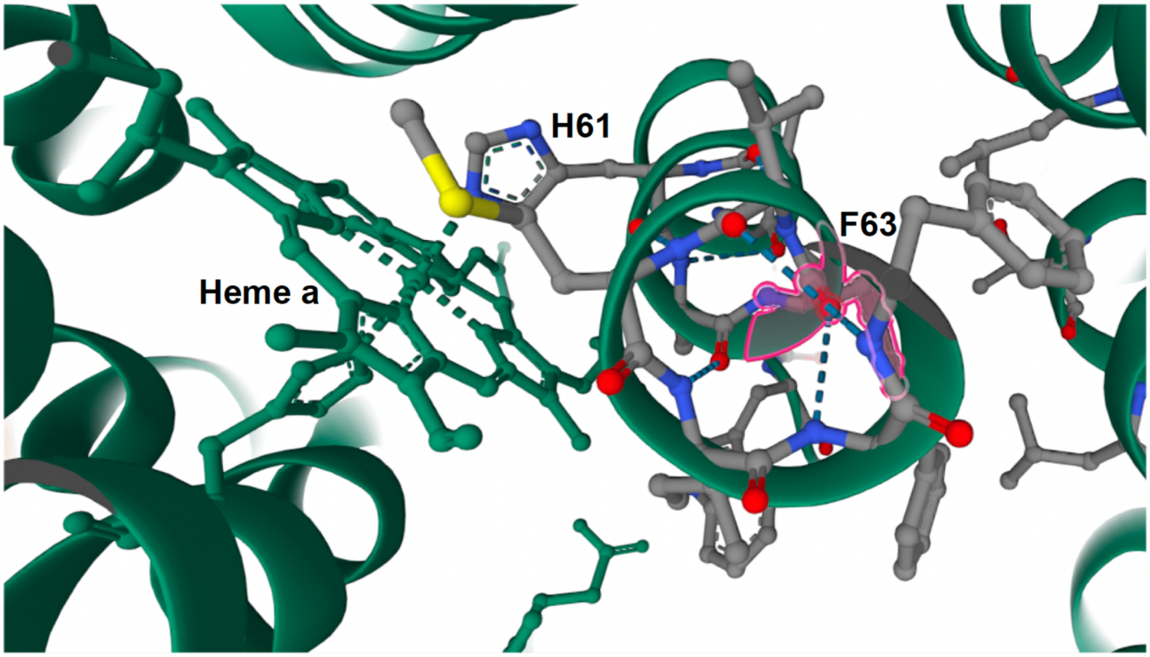
H61
F63
Heme a

**Figure S6. Comparison of the tri-dimensional structure of ATP6 A177T with the reference protein.** The reference protein is shown in dark blue and the variant in light blue. The A177 residue appears in red and the T177 variant residue in yellow. Alphafold was used to create the figure.

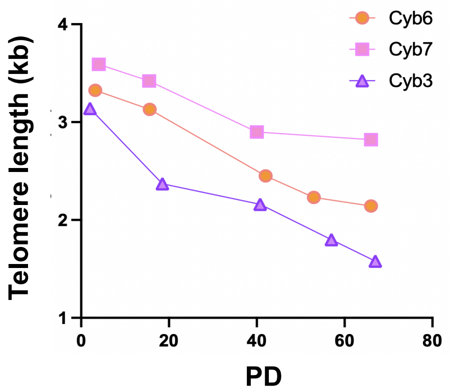
Cyb6
Cyb7
Cyb3
Telomere length (kb)
4
3
2
1
0
20
40
60
80
PD

**Figure S7. Longitudinal monitoring of telomere length in cultured Cyb3, Cyb6, and Cyb7 cells.** Following successive cell passaging, telomere length was quantified via TRF analysis at the designated population doubling (PD) intervals.

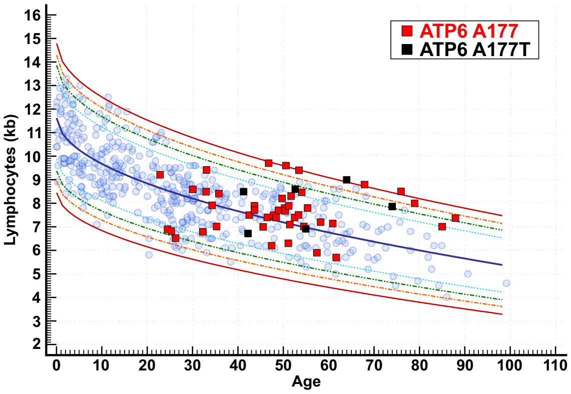

ATP6 A177
ATP6 A177T
Lymphocytes (kb)
Age

**Figure S8. Telomere length in the lymphocytes of donors with either m.G9055 (ATP6 A177) or m.G9055>A (ATP6 A177T).** Red squares: ATP6 A177; black squares: ATP6 A177T.The P50 curve is shown in dark blue.
